# Multiple prebiotic metals mediate translation

**DOI:** 10.1101/256958

**Authors:** Marcus S. Bray, Timothy K. Lenz, Jay William Haynes, Jessica C. Bowman, Anton S. Petrov, Amit R. Reddi, Nicholas V. Hud, Loren Dean Williams, Jennifer B. Glass

## Abstract

Today, Mg^2+^ is an essential cofactor with diverse structural and functional roles in life’s oldest macromolecular machine, the translation system. We tested whether ancient Earth conditions (low O_2_, high Fe^2+^, high Mn^2+^) can revert the ribosome to a functional ancestral state. First, SHAPE (Selective 2’ <u>H</u>ydroxyl <u>A</u>cylation analyzed by <u>P</u>rimer <u>E</u>xtension) was used to compare the effect of Mg^2+^, Fe^2+^, and Mn^2+^ on the tertiary structure of rRNA. Then, we used *in vitro* translation reactions to test whether Fe^2+^ or Mn^2+^ could mediate protein production, and quantified ribosomal metal content. We found that: (i) Mg^2+^, Fe^2+^, and Mn^2+^ had strikingly similar effects on rRNA folding; (ii) Fe^2+^ and Mn^2+^ can replace Mg^2+^ as the dominant divalent cation during translation of mRNA to functional protein; (iii) Fe and Mn associate extensively with the ribosome. Given that the translation system originated and matured when Fe^2+^ and Mn^2+^ were abundant, these findings suggest that Fe^2+^ and Mn^2+^ played a role in early ribosomal evolution.

**SIGNIFICANCE:** Ribosomes are found in every living organism where they are responsible for the translation of messenger RNA into protein. The ribosome’s centrality to cell function is underscored by its evolutionary conservation; the core structure has changed little since its inception ~4 billion years ago when ecosystems were anoxic and metal-rich. The ribosome is a model system for the study of bioinorganic chemistry, owing to the many highly coordinated divalent metal cations that are essential to its function. We studied the structure, function, and cation content of the ribosome under early Earth conditions (low O_2_, high Fe^2+^, high Mn^2+^). Our results expand the roles of Fe^2+^ and Mn^2+^ in ancient and extant biochemistry as cofactors for ribosomal structure and function.

---

Life arose around 4 billion years ago on an anoxic Earth with abundant soluble Fe^2+^ and Mn^2+^ (1–5). Biochemistry had access to vast quantities of these metals for over a billion years before biological O_2_ production was sufficient to oxidize and precipitate them. The pervasive use of these “prebiotic” metals in extant biochemistry, despite current barriers to their biological acquisition, likely stems from their importance in the evolution of the early biochemical systems.

The translation system, which synthesizes all coded protein (6, 7), originated and matured during the Archean Eon (4-2.5 Ga) in low-O_2_, high-Fe^2+^, and high-Mn^2+^ conditions (8). The common core of the ribosome, and many other aspects of the translation system, have remained essentially frozen since the last universal common ancestor (9). In extant biochemistry, Mg^2+^ ions are essential for both structure and function of the ribosome (10) and other enzymes involved in translation (11). In ribosomes, Mg^2+^ ions engaged in a variety of structural roles (Table 1), including in Mg^2+^-rRNA clamps (12, 13) (Fig. 1a), in dinuclear microclusters that frame the peptidyl transferase center (PTC) (13) (Fig. 1b), and at the small subunit-large subunit (SSU-LSU) interface (14) (Fig. 1c). Functional Mg^2+^ ions stabilize a critical bend in mRNA between the P-site and A-site codons (15) (Fig. 1d), and mediate rRNA-tRNA and rRNA-mRNA interactions (16) (Fig. 1e, f). Mg^2+^ ions also interact with some rProteins (17). Additionally, accessory enzymes needed for translation – aminoacyl tRNA synthetases, methionyl-tRNA transformylase, creatine kinase, myokinase, and nucleoside-diphosphate kinase – require Mg^2+^ ions as cofactors (Table 1).

**Table 1.** Structural and functional roles for select divalent cations in the translation system. All biomolecules in the table have been shown to require Mg^2+^ and may also be active with Fe^2+^ or Mn^2+^. “n.a.” indicates that data are not available.

| Translation system component(s) | Location of divalent ion | Role of divalent cation | Optimal $[Mg^{2+}]$ (mM) |
| --- | --- | --- | --- |
| **Ribosome** |  |  |  |
| LSU/SSU | $M^{2+}$-rRNA clamps (12) | Mediates and maintains folding/structure of rRNAs | ~10 (34) |
| LSU | Dinuclear microclusters (13) | Frames peptidyl transferase center (PTC) | ~10 (34) |
| LSU/SSU | LSU/SSU interface (27) | Mediates docking of mRNA to SSU and association of SSU with LSU | ~10 (34) |
| SSU/mRNA | Critical bend in mRNA between the P-site and A-site codons (16, 54) | Maintains correct reading frame mRNA | ~10 (34) |
| A-site tRNA/<br>P-site tRNA | tRNA-tRNA interface (27) | Stabilize tRNAs in the PTC | ~10 (34) |
| LSU/tRNA | rRNA-tRNA interface (27) | Stabilize rRNA-tRNA in the PTC | ~10 (34) |
| **Auxiliary** |  |  |  |
| EF-Tu | GTP binding site (55) | Stabilizes the transition state | 5-15 (56) |
| EF-G | GTP binding site (57) | Stabilizes the transition state | n.a. |
| Aminoacyl-tRNA synthetases | ATP binding site (58) | Stabilizes the transition state | >1 (59) |
| Methionyl-tRNA transformylase | ATP binding site (60) | Stabilizes the transition state | 7 (60) |
| Creatine kinase | NTP binding site (61) | Stabilizes the transition state | ~5 (61) |
| Myokinase | Acceptor NDP binding site (62) | Stabilizes the transition state | ~3 (45) |
| Nucleoside-diphosphate kinase | NTP binding site (63) | Stabilizes the transition state | >1 (63) |
| Pyrophosphatase | Active site (64) | Stabilizes the transition state | >7 (65) |

**Fig. 1.**
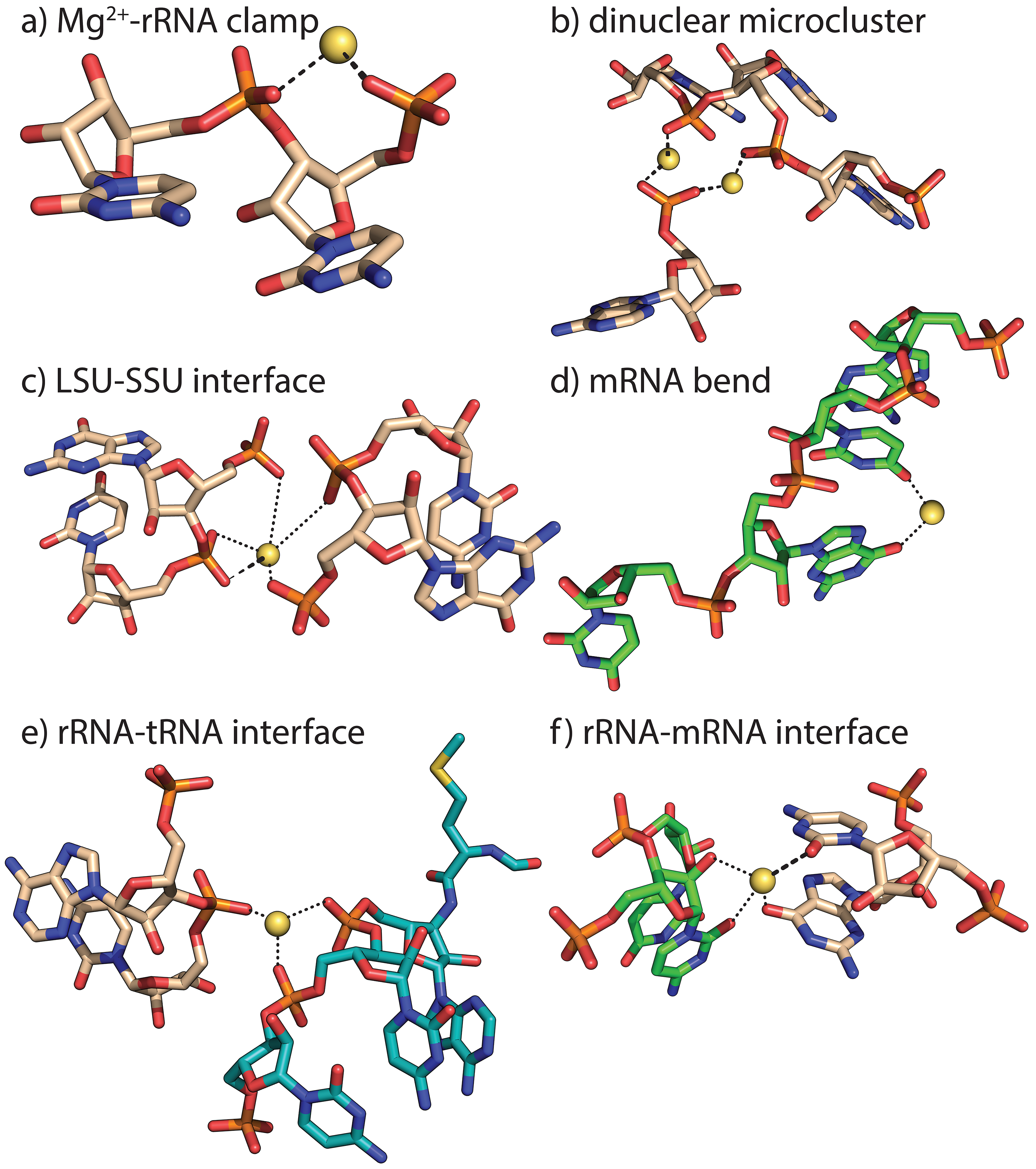
Divalent cations serve many structural and functional roles in the ribosome. Mg^2+^ ions: **a)** form bidentate clamps with adjacent phosphate groups of rRNA, **b)** form dinuclear microclusters that frame the rRNA of the PTC, **c)** stabilize the LSU-SSU interface, **d)** stabilize a functional kink in mRNA, **e)** stabilize association of tRNA (teal) with 23S rRNA (beige carbon atoms), and **f)** stabilize association of mRNA (green) with 16S rRNA (beige carbon atoms). Thick dashed lines are first shell RNA interactions of Mg^2+^. Dotted lines indicate second shell interactions. Images are of the *Thermus thermophilus* ribosome (PDB ID: 1VY4). This figure was generated with the program RiboVision (66).

Multiple types of cationic species can interact productively with RNAs in a variety of systems (18–20). Recent results support a model in which Fe^2+^ and Mn^2+^, along with Mg^2+^, were critical cofactors in ancient nucleic acid function (21). As predicted by this model, functional Mg^2+^-to-Fe^2+^ substitutions under anoxic conditions were experimentally verified to support RNA folding and catalysis by ribozymes (22, 23), a DNA polymerase, a DNA ligase, and an RNA polymerase (24). Functional Mg^2+^-to-Mn^2+^ substitution has long been known for DNA polymerases (24–26). For at least some nucleic acid processing enzymes, optimal activity is observed at lower concentrations of Fe^2+^ than Mg^2+^ (22, 24). Based on these previous results, we hypothesized that Fe^2+^ and Mn^2+^ could partially or fully replace Mg^2+^ during translation. In this study, we relocated the translation system to the low-O_2_, Fe^2+^-rich, or Mn^2+^-rich environment of its ancient roots, and compared its structure, function, and cation content under modern vs. ancient conditions.

## Results

### Fe^2+^ and Mn^2+^ fold LSU rRNA to a near-native state

To test whether Fe^2+^ or Mn^2+^ can substitute for Mg^2+^ in folding rRNA to a native-like state, we compared folding of LSU rRNA of the bacterial ribosome in the presence of Mg^2+^, Fe^2+^, or Mn^2+^ by SHAPE (<u>S</u>elective 2’-<u>H</u>ydroxyl <u>A</u>cylation analyzed by <u>P</u>rimer <u>E</u>xtension). SHAPE provides quantitative, nucleotide-resolution information about RNA flexibility, base pairing, and 3D structure, and has previously been used to monitor the influence of cations, small molecules, or proteins on RNA structure (27–32). We previously used SHAPE to show that the LSU rRNA adopts a near-native state in the presence of Mg^2+^, with the core inter-domain architecture of the assembled ribosome and residues positioned for interactions with rProteins (33). Here, SHAPE experiments were performed in an anoxic chamber to maintain the oxidation state of the metals and to prevent Fenton cleavage. The minimum concentration required to fully fold rRNA (10 mM Mg^2+^, 2.5 mM Fe^2+^, or 2.5 mM Mn^2+^) was used for all SHAPE experiments (**Datasets S1 and S2**).

Addition of Mg^2+^, Fe^2+^, or Mn^2+^ induced widespread structural changes in the LSU rRNA in the presence of Na^+^, as reflected in SHAPE profiles (see **Materials and Methods**) and displayed as ‘heat maps’ on the LSU rRNA secondary structure (Fig. 2;***SI Appendix*, Fig. S1**). Among the nucleotides forming the PTC, similar SHAPE profiles were obtained in the presence of Mg^2+^, Fe^2+^, or Mn^2+^ (***SI Appendix*, Fig. S1**). The ΔFe^2+^ and ΔMg^2+^ heat maps obtained for the entire 23S rRNA are nearly identical in most regions (Fig. 2d, e). As expected for conversion of secondary structure to fully folded tertiary structure, helices tended to be invariant, whereas loops and bulges were impacted by addition of Mg^2+^, Fe^2+^, or Mn^2+^. For the 23S rRNA, 86% of nucleotides (43/50) that exhibited a significant response (>0.3 SHAPE units) to Mg^2+^ also exhibited a similar trend with Fe^2+^. The greatest discrepancy between Fe^2+^ and Mg^2+^ was observed in the L11 binding domain (Fig. 2d, e).

**Fig. 2.**
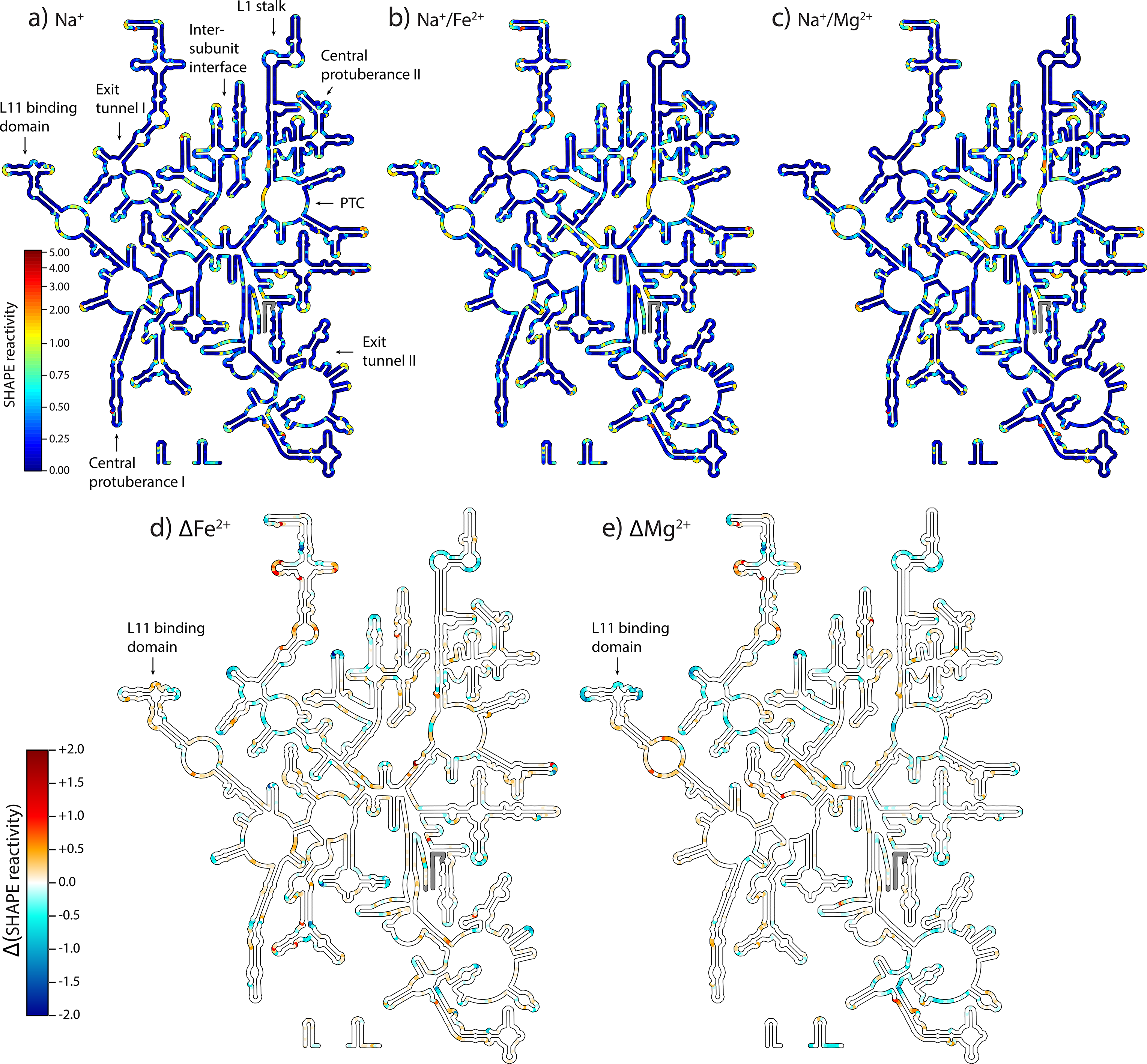
SHAPE reactivities mapped onto the *Thermus thermophilus* LSU rRNA secondary structure in **a)** Na^+^, **b)** Na^+^/Fe^2+^, or **c)** Na^+^/Mg^2+^. Key functional elements are labeled in panel a, and the scale in panel (a) applies to panels b and c. **d)** Fe^2+^-induced changes (ΔFe^2+^) in SHAPE reactivity calculated by subtracting Na^+^ data from Na^+^/Fe^2+^ data for each nucleotide, and **e)** Mg^2+^-induced changes (ΔMg^2+^) in SHAPE reactivity calculated by subtracting Na^+^ data from Na^+^/Mg^2+^ data for each nucleotide. The scale shown for panel d also applies to panel e. Positive values indicate increased SHAPE reactivity in presence of the divalent cation, while negative values denote decreased reactivity. Regions where data are not available (5’ and 3’ ends) are grey. These figures were generated with the program RiboVision (66). The L11 binding region, where the greatest discrepancy between Fe^2+^ and Mg^2+^ is observed, is indicated with an arrow.

### Fe^2+^ and Mn^2+^ mediate translation

Translation reactions were performed in an anoxic chamber in the presence of various cations and cation concentrations. Production of the protein dihydrofolate reductase (DHFR) from its mRNA was used to monitor translational activity. Protein synthesis was assayed by measuring the rate of NADPH oxidation by DHFR. These reactions were conducted in a small background of 2.5 mM Mg^2+^ (***SI Appendix*, Fig. S2a**). This background is below the requirement to support translation, consistent with previous findings that a minimum of ~5 mM Mg^2+^ is needed for assembly of mRNA onto the SSU (34, 35). As a control, we recapitulated the previously established Mg^2+^ dependence of the translation system, and then repeated the assay with Fe^2+^.

Activity of the translation system with variation in [Fe^2+^] closely tracks activity with variation in [Mg^2+^] (Fig. 3). Below 7.5 mM, total divalent cation concentration, minimal translation occurred with either Fe^2+^ or Mg^2+^, as expected (36). Activity peaked at 9.5 mM for both cations and decreased modestly beyond the optimum. At a given divalent cation concentration, Fe^2+^ supported around 50-80% of activity with Mg^2+^ (Fig. 4). This result was observed with translation reactions run for 15, 30, 45, 60, 90 and 120 min at the optimal divalent cation concentrations. Mn^2+^ also supported similar translation activity to Fe^2+^ at optimal divalent concentrations (***SI Appendix*, Fig. S3**). Along with Mg^2+^, Fe^2+^, and Mn^2+^ we investigated whether other divalent cations could support translation. No translation activity was detected with Co^2+^, Cu^2+^, or Zn^2+^ (***SI Appendix*, Fig. S3**).

**Fig. 3.**
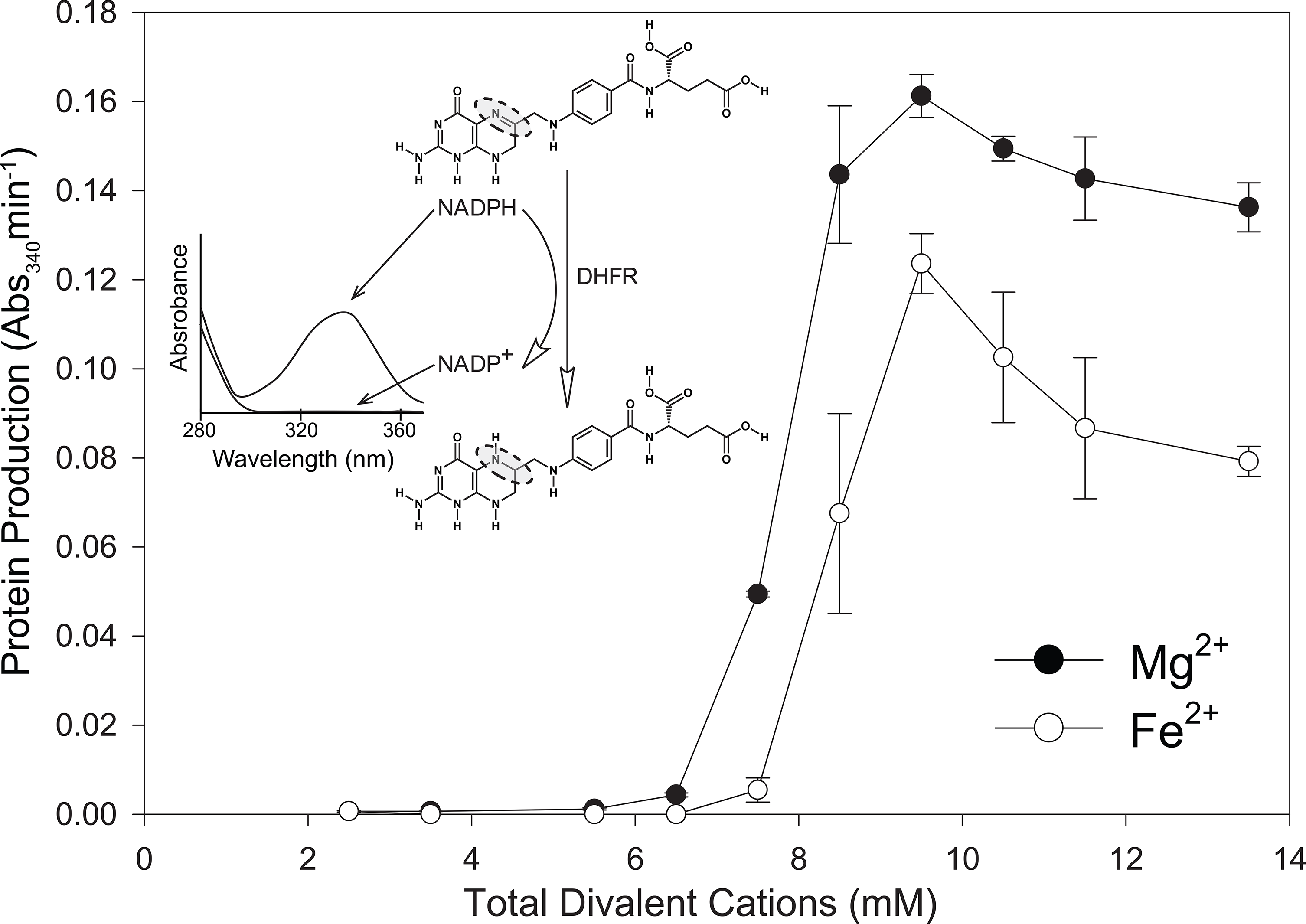
Mg^2+^ and Fe^2+^ stimulate translational activity over a range of concentrations. The activity of the translation product (dihydrofolate reductase, which catalyzes the oxidation of NADPH, with a maximum absorbance at 340 nm) was used as a proxy for protein production. Translation reactions were run for 120 minutes. All translation reactions contained 2.5 mM background Mg^2+^, to which varying amounts of additional Mg^2+^ or Fe^2+^ were added. The error bars for triplicate experiments (n=3) are plotted as the standard error of the mean.

**Fig. 4.**
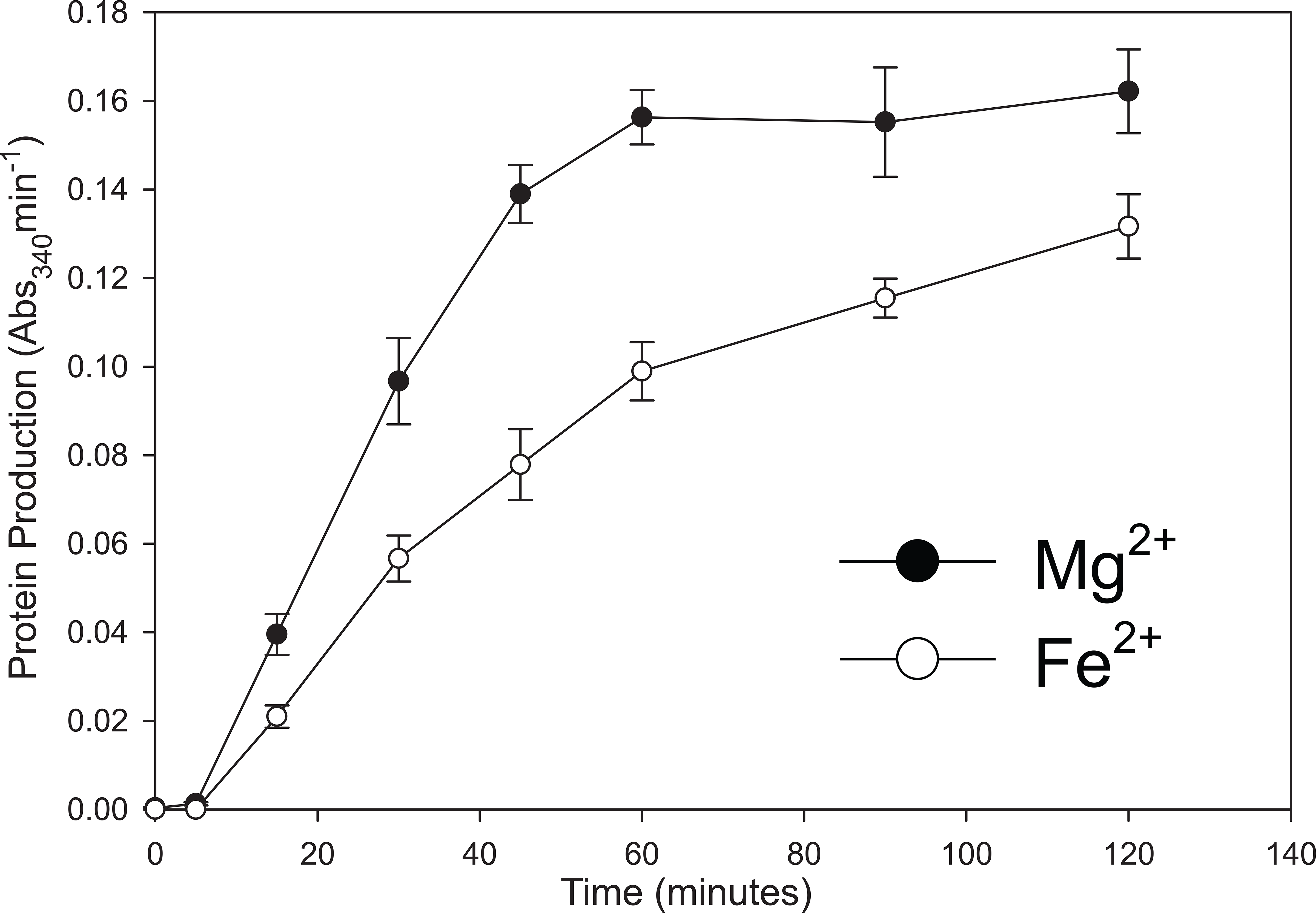
Fe^2+^ consistently supports 50-80% of the translational activity as Mg^2+^ when the translation experiments are run for 15-120 minutes. The activity of the translation product (dihydrofolate reductase, which catalyzes the oxidation of NADPH, with a maximum absorbance at 340 nm) was used as a proxy for protein production. All translation reactions contained 2.5 mM background Mg^2+^, to which 7 mM additional Mg^2+^ or Fe^2+^ were added, totaling to 9.5 mM divalent cation. The error bars for triplicate experiments (n=3) are plotted as the standard error of the mean.

To test whether alternative divalent cations could completely replace Mg^2+^ in translation, we decreased the background Mg^2+^ from 2.5 to 1 mM by thoroughly washing the ribosomes prior to translation reactions with 7-11 mM Fe^2+^ or Mn^2+^ (***SI Appendix*, Fig. S2b**). With 1 mM background Mg^2+^, Fe^2+^ supported 12-23% of the activity with Mg^2+^ over the concentrations tested, while Mn^2+^ supported 43-50% activity relative to Mg^2+^ (Fig. 5a). Washing the factor mix allowed us to decrease the background Mg^2+^ in translation reactions to ~4-6 μM (***SI Appendix*, Fig. S2c**). At this level, minimal protein production was observed with Fe^2+^, while Mn^2+^ supported 29-38% of the activity measured with Mg^2+^ (Fig. 5b).

**Fig. 5.**
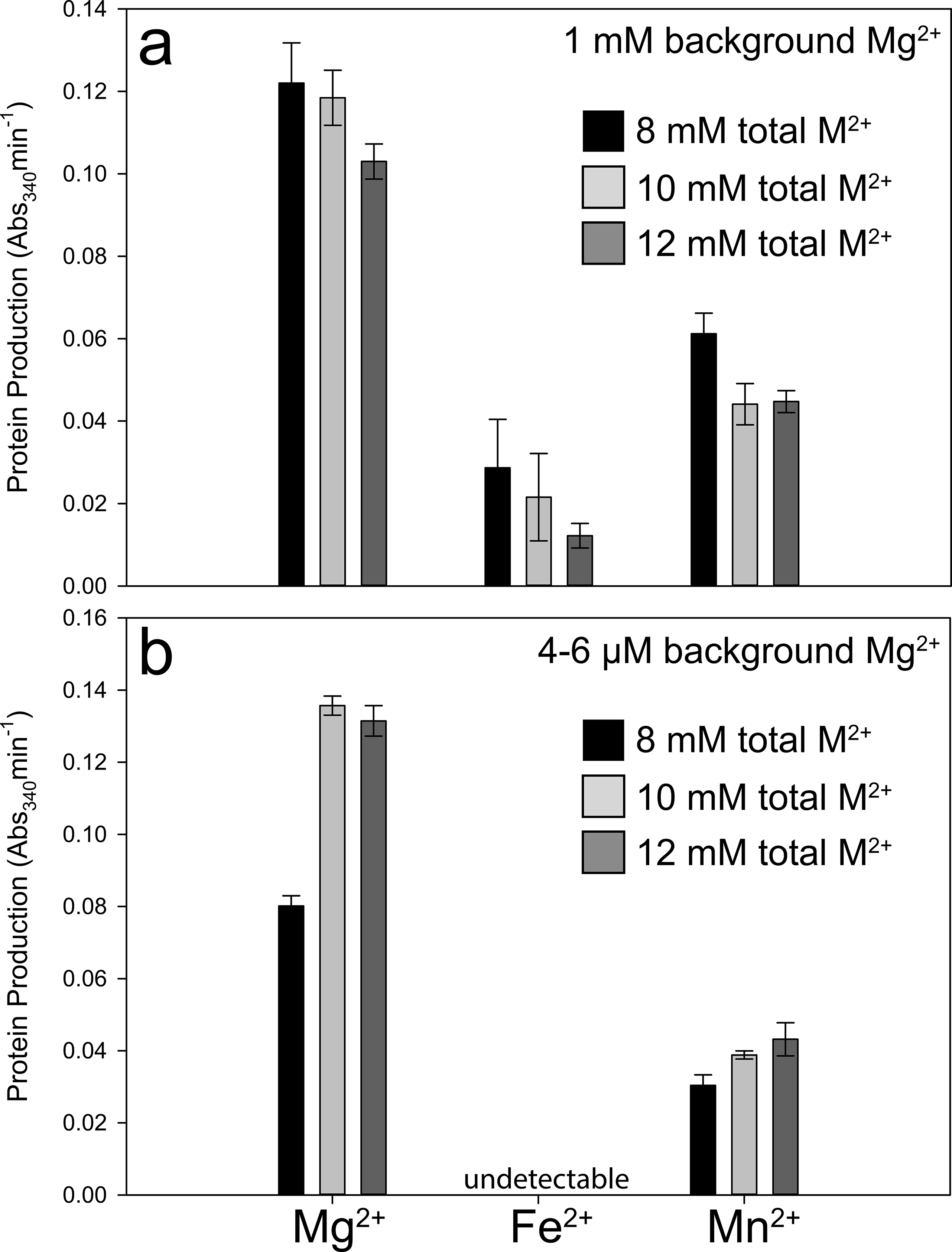
Mn^2+^ can support translation after removal of background Mg^2+^. **a)** Reactions prepared with washed *E. coli* ribosomes, reducing the background Mg^2+^ to 1 mM, to which 7, 9, or 11 mM additional Mg^2+^, Fe^2+^, or Mn^2+^ were added, totaling 8, 10, or 12 mM divalent cation (M^2+^). **b)** Reactions prepared using washed *E. coli* ribosomes and washed factor mix, which reduced the background Mg^2+^ to the low μM level, to which 8, 10, or 12 mM additional, Mg^2+^ Fe^2+^, or Mn^2+^ were added. The activity of the translation product (dihydrofolate reductase, which catalyzes the oxidation of NADPH, with a maximum absorbance at 340 nm) was used as a proxy for protein production. The error bars for triplicate experiments (n=3) are plotted as the standard error of the mean.

### Fe and Mn associate extensively with the ribosome

To experimentally confirm that Fe and Mn associate with the assembled ribosome, we analyzed the total Fe or Mn content of ribosomes after incubation in anoxic reaction buffer containing 7 mM Fe^2+^ or 7 mM Mn^2+^. Under the conditions of our translation reactions, 584 ± 9 Fe atoms or 507 ± 28 Mn atoms associate with each ribosome.

Finally, we computationally investigated whether Mg^2+^, Fe^2+^, and Mn^2+^ might be interchangeable during translation using quantum mechanical characterization of M^2+^-rRNA clamps (Fig. 1a; ***SI Appendix*, Fig. S4)**, which are abundant in the ribosome (12, 13). The geometries of Mg^2+^-rRNA, Fe^2+^-rRNA and Mn^2+^-rRNA clamps, are nearly identical (***SI Appendix*, Table S1-S3**). However, due to the accessibility of their d-orbitals, more charge is transferred to Fe^2+^ or Mn^2+^ (***SI Appendix*, Table S2**) than to Mg^2+^ (***SI Appendix*, Table S3)**. The effect of the modestly greater radius of Mn^2+^ (***SI Appendix*, Table S1)** is offset by d-orbital charge transfer (***SI Appendix*, Table S2)**, leading to elevated stability of Fe^2+^-rRNA clamp over the Mn^2+^-rRNA clamp (***SI Appendix*, Table S3)**.

## Discussion

In this study, we successfully replaced ribosomal Mg^2+^ with Fe^2+^ or Mn^2+^ under conditions mimicking the anoxic Archean Earth. Previously, the only divalent cation known to mediate rRNA folding and function was Mg^2+^. We found that isolated rRNA folds to essentially the same global state (37, 38) with Mg^2+^, Fe^2+^, or Mn^2+^ under anoxia. We then revealed that Fe^2+^ or Mn^2+^ can serve as the dominant divalent cation during translation. Mg^2+^ at 2.5 mM was insufficient to mediate protein synthesis; 5 mM additional Mg^2+^, Fe^2+^, or Mn^2+^ restored translational activity. These findings suggest that functional substitutions of Mn^2+^ or Fe^2+^ for Mg^2+^ can occur in large ribozymes, similar to previous reports for protein enzymes and small ribozymes (24–26, 39, 40). Near-complete removal of Mg^2+^ prevented Fe^2+^-mediated translation and partially inhibited Mn^2+^-mediated translation, suggesting that Mg^2+^ is optimal for some specific roles in the translation system. Regardless, the general effectiveness of Mn^2+^ or Fe^2+^ for Mg^2+^ substitutions in the translation system is astounding considering the enormous number of divalent cations associated with a given ribosome, and the broad scope of their structural and functional roles (10, 11) (Table 1, Fig. 1).

The observation that >500 Fe or Mn ions can associate with a bacterial ribosome is consistent with the number of Mg^2+^ ions observed by X-ray diffraction (100-1000 Mg^2+^ per ribosome (41)), and supports a model in which Fe^2+^ or Mn^2+^ has replaced Mg^2+^ as the dominant divalent cation in our experiments. The high capacity of ribosomes for Fe^2+^ and Mn^2+^ reflects all rRNA-associated divalent cations, including condensed, glassy and chelated divalent cations (42), and in addition, we presume that Fe^2+^ or Mn^2+^ can associate with a variety of rProteins, including those previously shown to bind Zn^2+^ (e.g. S2, S15, S16, S17, L2, L13, L31, L36 in *E. coli*) (43).

The differences in protein production observed among the three divalent cations likely arise from a variety of evolutionary and physiological factors. For instance, *E. coli* ribosomes may be evolutionarily adapted to Mg^2+^ instead of Fe^2+^ or Mn^2+^. The difference in translational activity between Mn^2+^ and Fe^2+^, particularly when background Mg^2+^ was removed, suggests that Mn^2+^ is more viable than Fe^2+^ upon full substitution. Mn^2+^/Mg^2+^ interchangeability may depend on relative stabilities of Mn^2+^ and Mg^2+^ in M^2+^-rRNA clamps (***SI Appendix*, Fig. S3**). Besides the ribosome, our translation reactions utilize many accessory proteins such as elongation factors and aminoacyl-tRNA synthetases that also have divalent cation requirements. Decreased activity of any of one these systems with Mn^2+^ and Fe^2+^ would cause a pinch point in an otherwise fully functional translation system. Indeed, the relative activity of myokinase and arginine t-RNA synthetase are both lower with Mn^2+^ or Fe^2+^ than with Mg^2+^ (44, 45).

While intracellular Mg^2+^ is around 10^−3^ M (46), specific physiological or environmental conditions can significantly elevate intracellular Fe^2+^ and Mn^2+^. Under oxidative stress, some microbes accumulate excess Mn^2+^. For example, radiation-tolerant *Deinococcus radiodurans* contains ~10 times higher Mn^2+^ than *E. coli* (~10^−5^ M Mn^2+^ (47, 48)). In the absence of O_2_, *E. coli* contains ~10 times higher labile Fe^2+^ (~10^−4^ M) than in the presence of O_2_ (~10^−5^ M (49)). Thus, it is possible that the absence of Fe^2+^ and Mn^2+^ in experimentally determined ribosomal structures is reflective of culturing, purification, or crystallization conditions (high O_2_, high Mg^2+^, low Fe^2+^, low Mn^2+^), and that other cations may also be present under diverse physiological conditions.

We have shown that the translation system functions with mixtures of divalent cations, which are variable during long-term evolutionary history and during short term changes in bioavailability and oxidative stress. When combined with previous results that DNA replication and transcription can be facilitated by Fe^2+^ and Mn^2+^ (18–20, 22–26, 39, 40), our findings that both Fe^2+^ and Mn^2+^ can mediate rRNA folding and translation of active protein has revealed that these “prebiotic” divalent metals can facilitate the entire central dogma of molecular biology (DNA➔RNA➔protein). These findings raise important questions about evolutionary and physiological roles for Fe^2+^ and Mn^2+^ in ancient and extant biological systems. Were Mg^2+^, Fe^2+^, and Mn^2+^ collaborators as cofactors on the ancient Earth, when Fe^2+^ and Mn^2+^ were more abundant (1–5), and Mg^2+^ was less abundant (2), than today? What was the role of Fe^2+^ and Mn^2+^ in the origin and early evolution of the translational system? Finally, what are the implications for ribosome-bound Fe^2+^ in oxidative damage and disease (50, 51)?

## Materials and Methods

### rRNA folding via SHAPE

SHAPE (28, 32, 33) was conducted on the ~2900 nt *Thermus thermophilus* 23 rRNA (LSU) in 250 mM monovalent cation (Na^+^ or K^+^) to favor formation of secondary structure, and in 250 mM Na^+^ or K^+^ plus various divalent cation (10 mM MgCl_2_, 2.5 mM FeCl_2_, or 2.5 mM MnCl_2_) to favor tertiary interactions. These divalent cation concentrations are sufficient to fold rRNA. To keep rRNA samples from O_2_, solutions of rRNA alone or 200 mM NaOAc or KOAc plus 50 mM Na-HEPES (pH 8) or K-HEPES (pH 8) and divalent cations were lyophilized and transferred into an anoxic chamber with a 98% Ar and 2% H_2_ atmosphere. The rRNA solutions were rehydrated with nuclease-free, degassed water, and added to the dried salts to achieve the appropriate concentrations. After rRNA modification reactions, divalent cations were removed by chelating beads. Samples were removed from the anoxic chamber before reverse transcription and analysis by capillary electrophoresis as in ref. (33). Essentially identical SHAPE profiles were observed with Na^+^ or K^+^ alone (***SI Appendix*, Fig. S1**), as previously described (32, 52); and for monovalent cations in combination Mg^2+^, Fe^2+^, or Mn^2+^ (Fig. 2; ***SI Appendix*, Fig. S1**). Nucleotides were classified as exhibiting a significant change in SHAPE reactivity if the difference between the initial reactivity (in Na^+^) and final reactivity (in Na^+^/Mg^2+^, Na^+^/Fe^2+^, or Na/Mn^2+^) was >0.3 SHAPE units. To compare the Mg^2+^-, Fe^2+^-, and Mn^2+^ - responsiveness of specific nucleotides, we binned nucleotides into three categories (increased, decreased, or little/no change) based on their general SHAPE reactivity response to each divalent cation (SHAPE data are found in **Datasets S1 and S2).**

### *In vitro* translation

Each 30 μL reaction contained 2 μM (4.5 μL of 13.3 μM stock) *E. coli* ribosomes in 10 mM Mg^2+^ (New England Biolabs, Ipswich MA, USA; catalog # P0763S), 3 μL factor mix (with RNA polymerase, and transcription/translation factors in 10 mM Mg^2+^) from the PURExpress^®^ Δ Ribosome Kit (New England Biolabs E3313S), 0.1 mM amino acid mix (Promega, Madison WI, USA; catalog # L4461), and 0.2 mM tRNAs from *E. coli* MRE 600 (Sigma-Aldrich, St. Louis MO, USA; product # TRNAMRE-RO). Thus, a total of 2.5 mM “background” Mg^2+^ was present in each reaction (***SI Appendix*, Fig. S2a**). To remove the background Mg^2+^, we exchanged the buffer of the ribosome and factor mix using centrifugal filter units. Thirty microliters of either ribosome solution or factor mix was added to an Amicon Ultra 0.5 mL centrifugal filter (Millipore-Sigma), followed by 450 μL of divalent-free buffer (20 mM HEPES pH 7.6, 30 mM KCl, and 7 mM β-mercaptoethanol). Samples where spun at 14,000 × g at 4C until the minimum sample volume (~15 μL) was reached. The samples were resuspended in 450 μL of divalent-free buffer and centrifugation was repeated. The samples were then transferred to new tubes and 15 μL of divalent-free buffer was added to bring the volume to 30 μL. This process decreased Mg^2+^ concentrations in the ribosome and factor mix from 10 mM to 10-30 μM Mg^2+^, resulting in 4-6 μM Mg^2+^ in each reaction (***SI Appendix*, Figs. S2b, S2c)**.

### Translation experimental conditions

All reactions (30 μL total volume) were assembled and incubated in an anoxic chamber. Divalent cation salts (MgCl_2_, FeCl_2_, MnCl_2_, Zn(OAc)_2_, CoCl_2_, CuSO_4_) were added to 7 mM final concentration, with the exception of MgCl_2_ and FeCl_2_, which were tested over a range of concentrations (1, 3, 4, 5, 6, 7, 8, 9, and 11 mM; ***SI Appendix*, Fig. S2**). Solutions were clear, with no indication of metal precipitate, suggesting that reduced, divalent metals cations were the primary chemical species. All experiments were assembled in the following order: dihydrofolate reductase (DHFR) mRNA (~5 μg per 30 μL reaction), factor mix, ribosomes, amino acids, tRNA, nuclease-free H_2_O, reaction buffer (see ***SI Appendix*** for details on mRNA template and reaction buffer recipe). Changing the order of reactant addition did not affect translational activity. Reactions were run in triplicate on a 37°C heat block for up to 120 minutes. Reactions were quenched on ice and stored on ice until they were assayed for protein synthesis.

### Protein activity assay

Protein synthesis was measured using a DHFR assay kit (Sigma-Aldrich product # CS0340), which measures the oxidation of NADPH (60 mM) to NADP^+^ by dihydrofolic acid (51 μM). Assays were performed by adding 5 μL of protein synthesis reaction to 995 μL of 1x assay buffer. The NADPH absorbance peak at 340 nm (Abs_340_) was measured in 15 s intervals over 2.5 min. The slope of the linear regression of Abs_340_ vs. time was used to determine protein activity (Abs_340_ min^−1^). Different counter ions (Cl^−^, CH_3_COO^−^, SO_4_^2-^) had no effect on protein synthesis from mRNA. To our knowledge, no dependence on, nor inhibitory effect of Mg^2+^ or Fe^2+^ exists for DHFR. We confirmed this by varying the metal concentrations in our assay reaction, which had no effect on DHFR activity.

### Ribosome metal content

The Fe and Mn content of *E. coli* ribosomes was measured by total reflection X-ray fluorescence (TRXF) spectroscopy after the ribosomes were incubated in 7 mM FeCl_2_ or 7 mM MnCl_2_. See ***SI Appendix*** for additional details.

### Quantum mechanical calculations

The atomic coordinates of a Mg^2+^-rRNA clamp were initially extracted from the X-ray structure of the *Haloarcula marismortui* LSU (PDB 1JJ2) (53). The free 5′ and 3′ termini of the phosphate groups were capped with methyl groups in lieu of the remainder of the RNA polymer, and hydrogen atoms were added, where appropriate (***SI Appendix*, Fig. S4**). Additional details on calculations adapted from previous publications (12, 22) are described in ***SI Appendix***.

## ACKNOWLEDGMENTS

This research was supported by the National Aeronautics and Space Administration grants NNX14AJ87G, NNX16AJ28G, and NNX16AJ29G. We acknowledge helpful discussions with Corinna Tuckey of New England Biolabs, and with Eric B. O’Neill and Claudia Montllor Albalate of Georgia Institute of Technology. We thank Michael Goodisman for access to a capillary electrophoresis instrument.

## Author contributions

MSB, TKL, JCB, ARR, NVH, LDW, and JBG conceived and designed the experiments. MSB, TKL, and JWH collected data and performed analysis. ASP contributed to analysis. MSB, TKL, LDW, and JBG wrote the paper with editorial input from all authors.

